## Supplementary Material for "Annexin- and calcium-regulated priming of legume root cells for endosymbiotic infection"

**This PDF includes:**

Supplementary Figures S1 to S16

Supplementary Tables S1 and S2

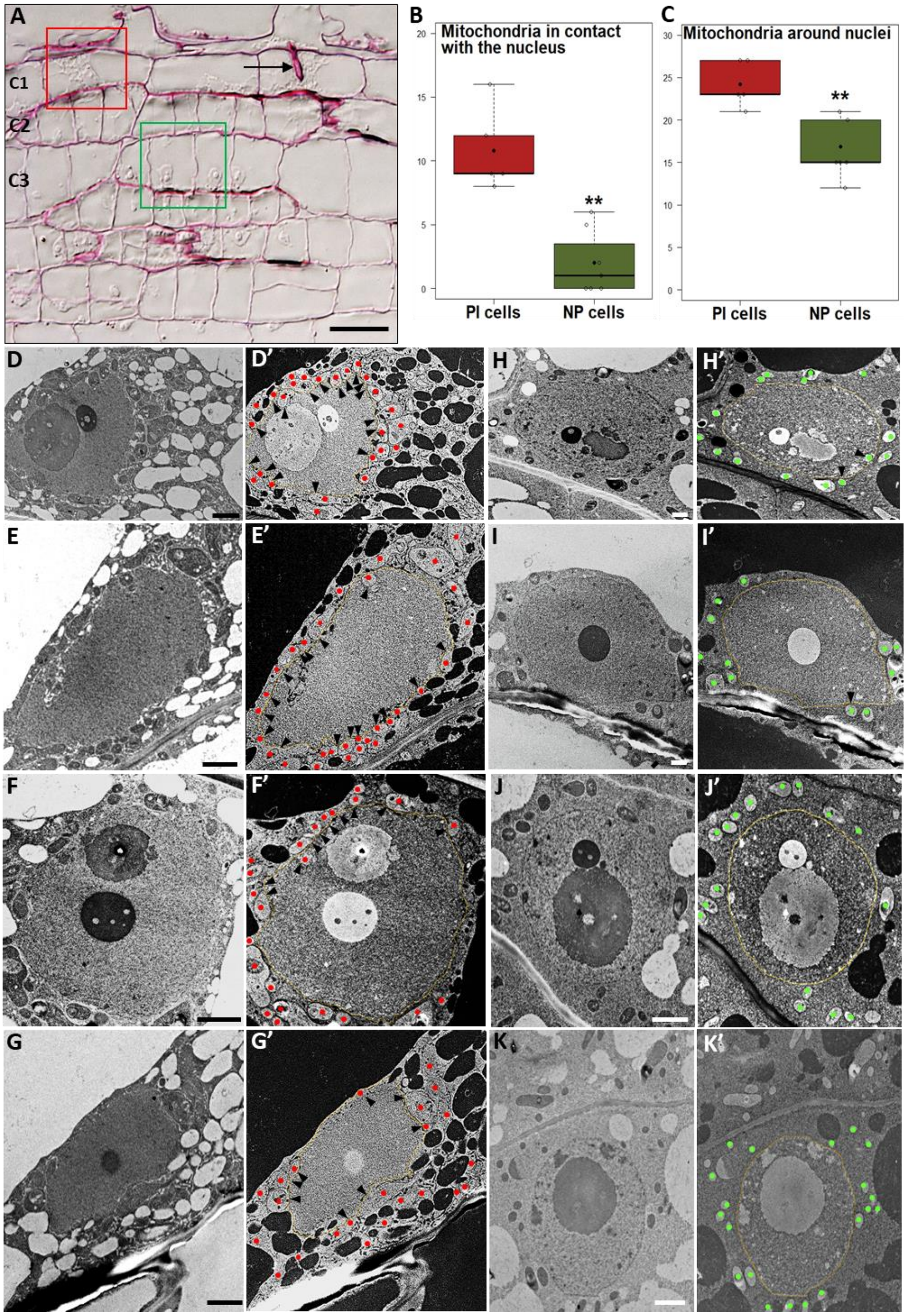


**Supplementary Fig. S1. Quantification of mitochondria in the vicinity of nuclei in rhizobia pre-infection-primed (PI) and dividing nodule primordia (NP) cells (related to Fig.1)**.

**(A)** A representative image of a 1 µm longitudinal section of an early rhizobia root colonization site (5 dpi) shows a primed cortical C1 cell before (red square) and during IT penetration (arrow). (**B**-**C**) The number of mitochondria in close contact with the nuclear envelope (**B**) or within a 2 µm radius around the nucleus were quantified in images of 80 nm-sections of C1-C2 cells in a pre-infection state (PI cells, as in the red box plot) or in dividing C3-C5 nodule primordia cells (NP cells, as in the green boxplots). Measurements were performed on n=8 (PI cells) and n=7 (NP cells) sections derived from 5-7 independent sites from 2 independent experiments. Box plots represent the distribution of individual values (indicated by open circles). Median (central line) and mean (solid black circle) values are indicated. A two-tailed Mann Whitney test (**B**) and a two-tailed t-test (**C**) were performed in R (asterisks indicate statistical difference; **p<0.01). (**D**-**K**) Representative images of PI cells (**D**-**K**) or dividing NP cells (**H**-**K**), used to quantify mitochondria in **B**-**C**. (**D'**-**K'**) LUT inverted images of **D**-**K.** Mitochondria are indicated by red dots in PI cells and by green dots in NP cells, black arrowheads indicate mitochondria in contact with the nucleus and yellow dotted lines underline nuclear contours. Scale bars: **A** = 50 µm, **D**-**K** = 2 µm.

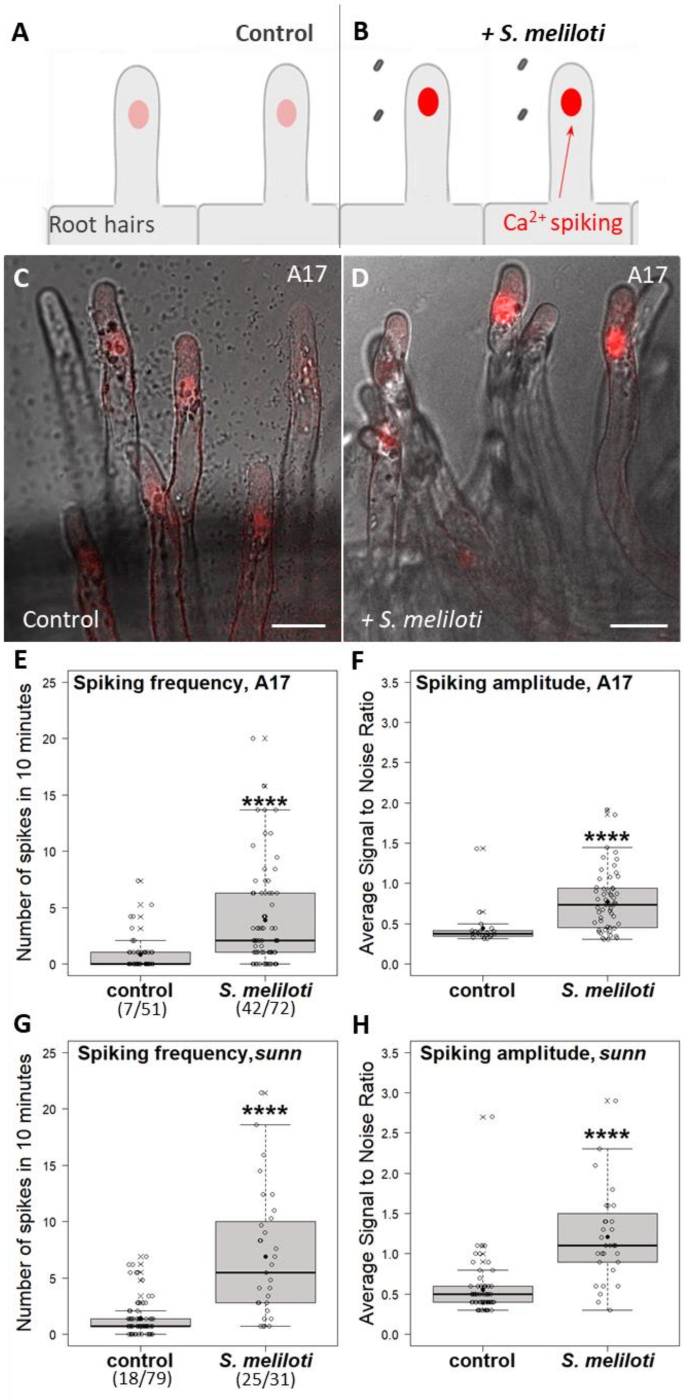


**Supplementary Fig. S2. Root hair $Ca^{2+}$ spiking responses to rhizobial inoculation in A17 and *sunn* (related to Fig. 2).**

(**A**-**D**) Schematic illustration (**A**-**B**) and representative confocal images (**C**-**D**) of nuclear NR-GECO1 fluorescence (in red) in Control, non-inoculated (**A**, **C**) and non-infected A17 root hairs of the nodulation-susceptible root zone 1 dpi with *S. meliloti* (**B**, **D**). Images in **C**-**D** are part of Supplementary movies M1 and M2, respectively. (**E**-**H**) Spiking frequency (**E**, **G**), expressed as number of spikes in 10 minutes per nucleus and spiking amplitude (**F**, **H**), expressed as average signal-to-noise ratio (SNR, cf. Methods section) of spikes per nucleus, in A17 (**E**, **F**) and *sunn* (**G**, **H**). Box plots in **E**-**G** represent the distribution of individual values from (**E**) Control (n = 51) and inoculated (+ *S. meliloti*, n=72) A17 root hairs, and (**G**) Control (n = 79) and inoculated (+

*S. meliloti*, n=31) *sunn* root hairs. Parentheses indicate number of root hairs with spiking/total number of root hairs. Root hairs are counted as spiking when showing more than 2 peaks in 10 min. Box plots in **F**-**H** represent the distribution of average values for root hairs showing at least 1 peak i.e. from (**F**) Control (n = 20) and inoculated (+ *S. meliloti*, n=56) A17 root hairs and (**H**) Control (n = 63) and inoculated (+ *S. meliloti*, n=31) *sunn* root hairs. Individual datapoints (open circles), median (central line), mean (solid black circle) and outliers (cross) are indicated. Data was acquired from three independent biological experiments for A17 (1dpi) and three independent biological experiments for *sunn* (2-4dpi). Two-tailed Mann-Whitney tests were performed for values in **E**-**H** (asterisks indicate significant differences, ****p<0.0001). Scale bars in **C**-**D** = 20 µm.

**Supplementary Fig. S3. $Ca^{2+}$ spiking responses restricted to outer cortical cells near rhizobial root hair infection sites in *sunn* (related to Fig. 3).**

$Ca^{2+}$ spiking responses were recorded in primed outer cortical cells neighbour to an infected root hair site in *M. truncatula sunn* roots expressing NR-GECO1 $Ca^{2+}$ sensor 2-7 dpi with CFP-expressing *S. meliloti* (in magenta). Representative images and $Ca^{2+}$ spiking traces collected from two independent rhizobia infection sites are shown (**A**-**D** and **E**-**H**, respectively). (**A**, **E**) Root hair (RH) infection threads (ITs, arrows) and the position of the root hair epidermal body (dotted white line) are indicated. (**B**, **F**) Outer cortical cells located in the area of the infected root hairs. The positions of the infected root hair(s) epidermal bodies are indicated as grey dotted lines. Outer cortical nuclei (C1a-m in **B**, C1a-g in **F**) analysed for $Ca^{2+}$ spiking (**D**, **H**) are indicated (round/oval dotted shapes). Non-labelled nuclei are either epidermal nuclei or those that were not in focus during the NR-GECO1 fluorescence acquisition. (**D**, **H**) $Ca^{2+}$ spiking traces from the nuclei of the 13 (**D**) and 7 outer cortical cells (**E**) labelled in **B** and **F**. At both infection sites, only a few cortical cells situated near the epidermal bodies of the infected root hair(s) are spiking at low frequency (cells C1b-d and C1h in **B**, cells C1b-d in **F**, their contours are indicated). Among them, C1c and C1b cells in **B**-**C** and **F**-**G,** respectively, are those that host an infection thread at a later timepoint (arrowheads in **C**, **G**). $Ca^{2+}$ spiking traces in **D**, **H** are expressed as signal-to-noise ratio (SNR). Images in **A**-**C** and **E**-**G** are maximal z-projections of sub-stacks. Scale bars = 40 µm.

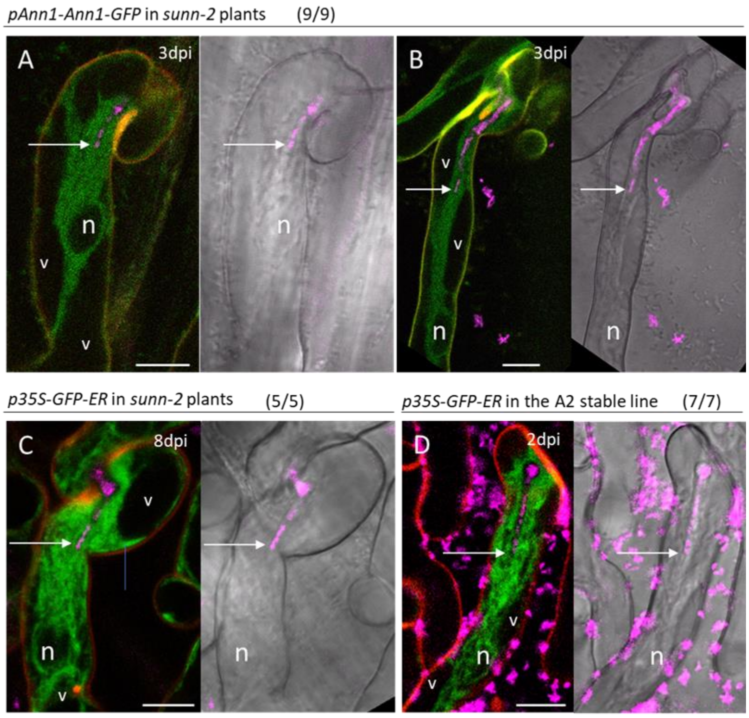


**Supplementary Fig. S4. GFP-ER marker and MtAnn1-GFP label root hair cytoplasmic bridges (related to Figs. 2-3).**

Root hairs undergoing infection were imaged 2 to 8 dpi with CFP-expressing *S. meliloti* (in magenta). Images were taken from root hairs of *M. truncatula sunn* composite plants expressing *pAnn1-Ann1-GFP* (**A**, **B**) or *p35S-GFP-ER* (**C**) or a stable line (referred to as A2) expressing *p35S-GFP-ER* (**D**). In these root hairs, elongating ITs are immersed in a cytoplasmic bridge connecting the IT to the nucleus (n). The cytoplasm bridges delimited by the vacuole (v, in black in the fluorescence images) are entirely and similarly labelled by MtAnn1-GFP (**A**, **B**) or GFP-ER (**C**, **D**). Arrows indicate the tip of the bacteria file within the growing infection thread. Images are merges of GFP (green), CFP (magenta) and autofluorescence (red) in left panels, or merged CFP and bright field in right panels. Parentheses (**A**-**D**) indicate the number of sites showing this pattern/total number of documented sites from 4 (**A**-**B**), 1 (**C**) and 3 (**D**) independent experiments, respectively. All sites were observed at a later time point to confirm elongation of the imaged ITs. Scale bars = 10 µm.

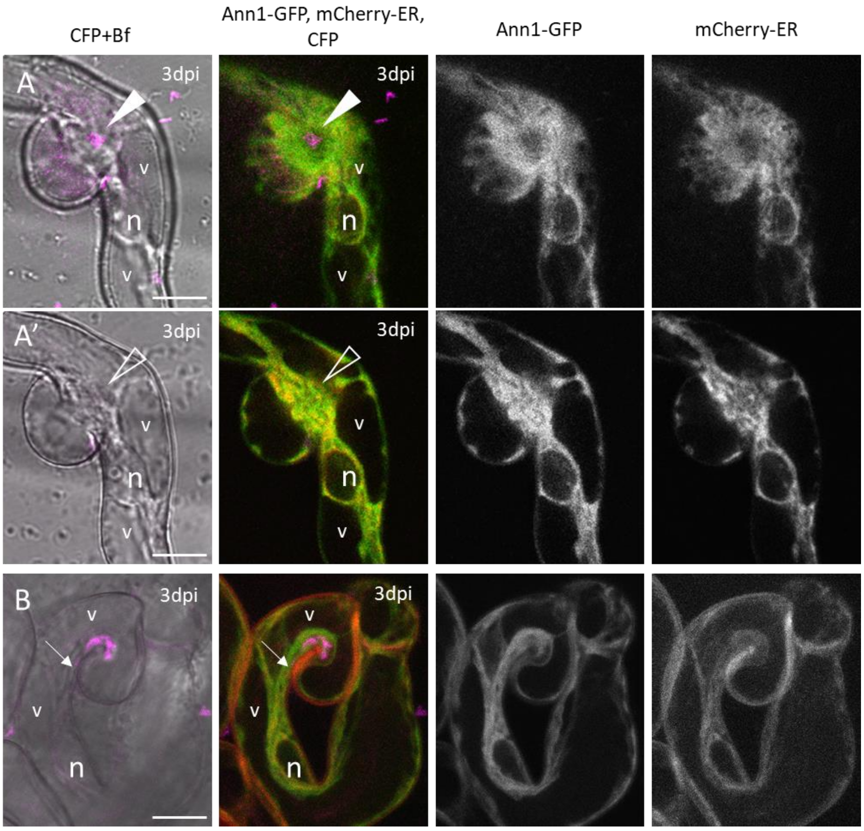


**Supplementary Fig. S5. MtAnn1-GFP and mCherry-ER both label cytoplasmic bridges in *M. truncatula* root hairs (related to Figs. 2-3).**

Composite *sunn* plants expressing both *pAnn1-Ann1-GFP* and *p35S-mCherry-ER* were inoculated with CFP-expressing *S. meliloti* (in magenta). Root hairs at early stages of infection (RHE and IT) were imaged at 3 dpi. Merged bright-field (Bf) and CFP (in magenta) are shown in the first column. Ann1-GFP and mCherry-ER fluorescences (in green and red, respectively) are merged in the second column, and showed separately (in grey) in the third and fourth columns as indicated. Colocalization of both markers in the second column is visualized in yellow. (**A**, **A'**) RHE stage, a curled root hair branch displaying an active, rhizobia-colonized infection chamber (**A**, arrowhead). **A'** shows a distinct focal plane of the same root hair as **A**, focused on the cytoplasmic bridge linking the infection chamber and the nucleus. This cytoplasmic bridge is highlighted similarly by MtAnn1-GFP and mCherry-ER. (**B**) IT stage, a curled RH hosting a developing IT (arrow). The cytoplasm in this RH, including the column linking the developing IT and the nucleus, is marked by MtAnn1-GFP and mCherry-ER alike. The site in **A**, **A'** is representative of 3 such sites in 2 independent experiments. **B** confirms the MtAnn1-GFP and GFP-ER patterns observed independently in other experiments. Images in **A** are from a single section, images in **A'** and **B** are maximal z-projections of 2 and 3 successive sections, respectively. Filled arrowhead, rhizobia in the infection chamber, open arrowhead, relative position of the infection chamber, arrow, tip of the rhizobia file within the infection thread, n, nucleus, v, vacuole. Scale bars = 10 µm.

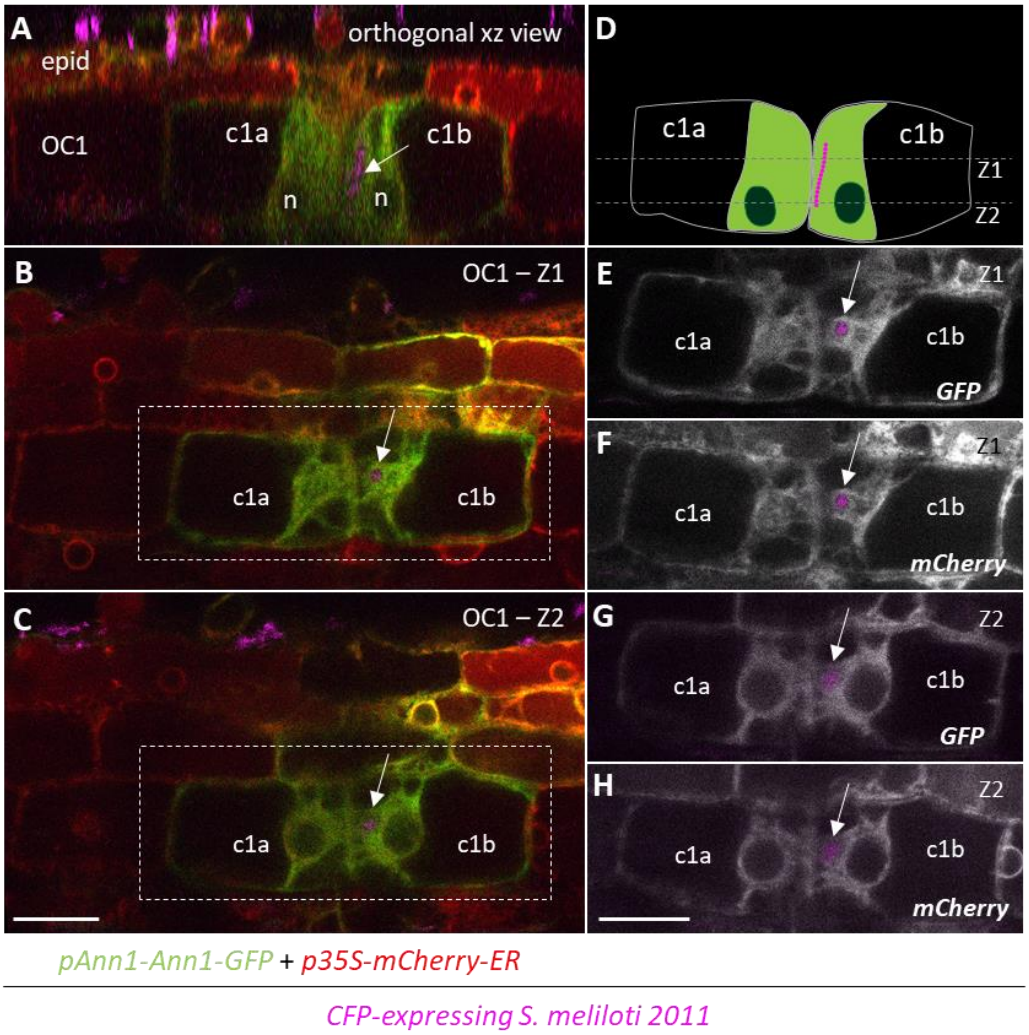


**Supplementary Fig. S6. ER marker and MtAnn1-GFP co-label cytoplasmic bridges in pre-infection primed cortical cells (related to Figs. 2-3).** Composite *sunn* plants co-expressing *pAnn1-Ann1-GFP* and *p35S-mCherry-ER* were inoculated with CFP-expressing *S. meliloti* (in magenta). Images in the left column (**A**-**C**) are merges of CFP (magenta), GFP (green), and mCherry (red), while images in the right column are merges of CFP (magenta) and either GFP (**E**, **G**) or mCherry (**F**, **H**) in grey. (**A**) Side view (or orthogonal xz view) of two outer cortical cells, c1a and c1b. These cells are adjacent to an infected root hair (not shown) and the c1a cell is preparing for infection while c1b is being infected as revealed by the presence of a segment of the infection thread (arrow). (**B**, **C**) show two distinct optical xy sections (Z1 and Z2) across the c1a and c1b cells, as indicated in the cartoon (**D**). (**E**-**H**) show separate GFP and mCherry channels for section Z1 (**E**, **F**) and Z2 (**G**, **H**), respectively. The typical cytoplasm organization in the activated c1a cell and infected c1b cell is similarly labelled by MtAnn1-GFP (**E**, **G**) and mCherry-ER (**F**, **H**). All images in **B**-**C** and **E**-**H** are single optical sections. Arrow, rhizobia in the infection thread, n, nucleus. Scale bars = 20 µm. These images are representative of 3 primed cell sites and 6 infected outer cortex cell sites acquired in 5 plants in 1 experiment.

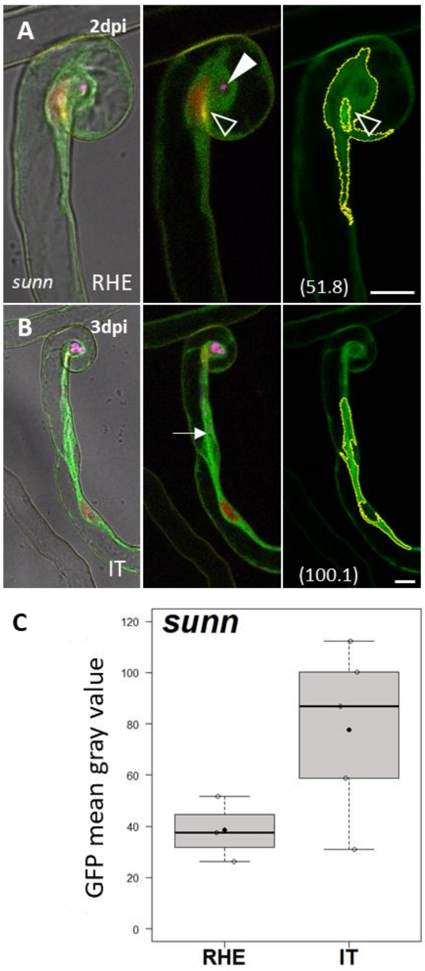


**Supplementary Fig. S7. MtAnn1-GFP signal intensity in RHE and IT root hair cells (related to Fig. 2).**

(**A**, **B**) MtAnn1-GFP fluorescence intensity was measured in root hairs at RHE (root hair with entrapped rhizobia) or IT (root hair with a growing infection thread) stages in *M. truncatula* *sunn*, representative images are shown in **A** (RHE) and **B** (**IT**). Root hairs were imaged in roots expressing MtAnn1-GFP (green) and NR-GECO1 (red), after inoculation with CFP-expressing *S. meliloti* (magenta). Merges of CFP, GFP and NR-GECO with (left panels) or without (central panels) bright field images are shown. Right panels show GFP fluorescence alone and the contours (yellow lines) of the areas (ROIs) used to measure mean grey fluorescence levels (numbers in parentheses) in the cytoplasm aggregated around the site of rhizobia entrapment (**A**, arrowhead) or the cytoplasmic bridge linking an elongating infection thread (**B**, arrow) and the nucleus. Note that in this site, a second, smaller ROI (open arrowhead), corresponding to the auto-fluorescent wall domain (or contact point) adjacent to infection chambers in curled root hairs[26] was subtracted from the analysis. Box plots in **C** represent the distribution of individual values from RHE (n = 3) and IT (n=5) *sunn* root hairs. Individual values (open circles), median (central line), mean (solid black circle) and outliers (crosses) are indicated. Arrowhead, site of rhizobia entrapment, open arrowhead, strongly auto-fluorescent wall contact point, arrow, infection thread. Scale bars = 10 µm.

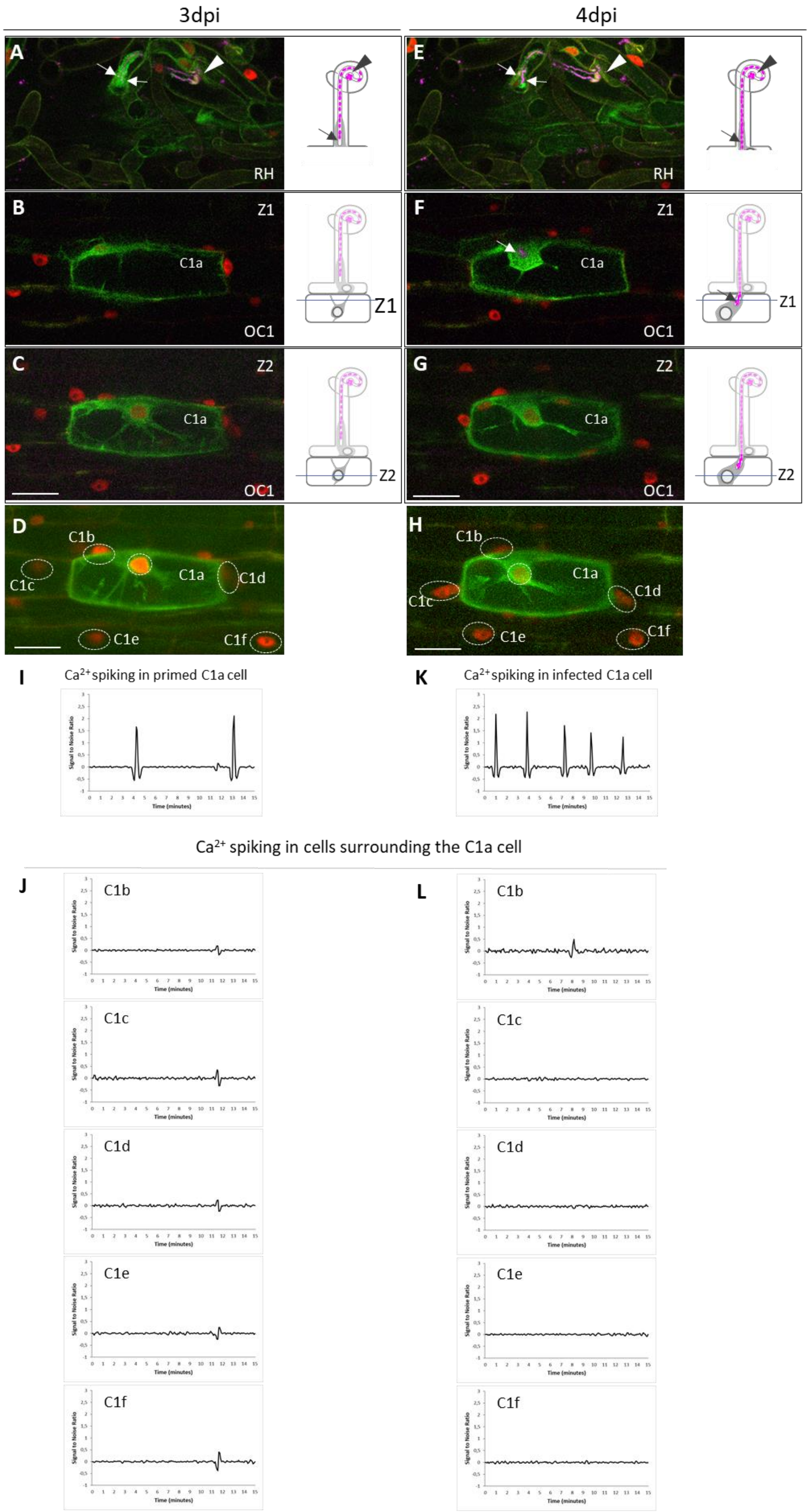


**Supplementary Fig. S8. MtAnn1-GFP dynamics and $Ca^{2+}$ spiking in primed and infected cortical cells in *sunn* (related to Fig. 3).**

(**A**-**L**) MtAnn1-GFP dynamics and $Ca^{2+}$ spiking in cortical cells of *sunn* expressing *pMtAnn1-MtAnn1-GFP* (green) and *NR-GECO1* (red) below an infected root hair site (**A**-**E**) at two successive stages, 3 dpi (cortical primed state **B**-**C**) and 4 dpi (cortical infection, **F**-**G**) with CFP-labelled *S. meliloti* (magenta). **A**-**C** and **E**-**G** associate confocal images (left panels) and cartoons (right panels) of the different cell layers and features of cells preparing for infection (**B**-**C**) or infected (**F**-**G**). (**D**, **H**) illustrate the regions of interest (ROIs, dotted oval shapes) used for $Ca^{2+}$ spiking analysis (**I**-**J**, **K**-**L**). (**B**-**D**, **F**-**H**) focus on the C1a outer cortical cell next to the infected root hair before (at 3dpi) and during (at 4dpi) rhizobia IT progression (**F**, arrow, Z1 section). Two virtual sections (Z1, Z2) across C1a illustrate cytoplasmic reorganisation in the pre-infection primed state (**B**-**C**) and later during infection (**F**-**G**). (**I**-**J**, **K**-**L**) show nuclear $Ca^{2+}$ traces from C1a cell when primed to be infected (3 dpi, **I**) or during infection (4 dpi, **K**) and from surrounding outer cortical cells C1b-f (**J**, **L**). Spiking frequency is higher in C1a during IT progression (**K**) than before (primed state, **I**). At both timepoints, only the MtAnn1-GFP-labelled C1a cell is spiking. Images in **A**, **C**, **E**, **G** are maximal z-projections of 12 (**A**, **E**, **G**) or 25 (**C**) successive sections, images in **B** and **F** are single confocal sections. Images in **D** and **H** are maximal projections of whole time series. The images are representative of 3 such sites in *sunn*. $Ca^{2+}$ traces are expressed as signal-to-noise ratio (SNR). Arrowhead, infection chamber, arrows, infection threads, RH, root hair, OC1, first outer cortical cell layer. Scale bars = 40 µm.

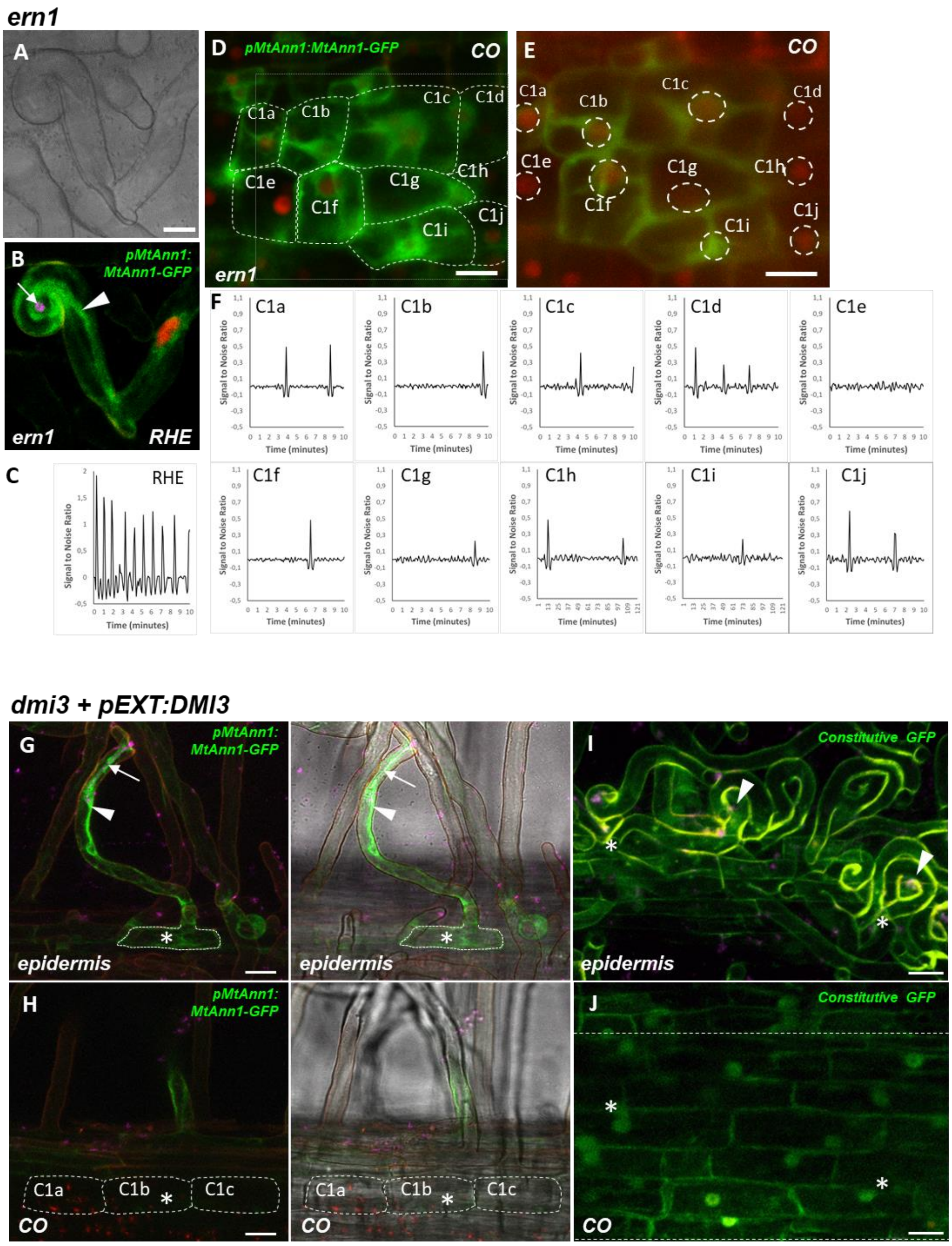


**Supplementary. Fig. S9. Infection priming in symbiotic defective mutants (related to Fig. 3).**

(**A**-**F**) The nuclear NR-GECO1 $Ca^{2+}$ sensor (red fluorescence) and the MtAnn1-GFP fluorescent fusion (green) were co-expressed in roots of the *ern1* mutant. (**A**-**B**) Bright-field and corresponding confocal images of an *ern1* root hair with entrapped rhizobia (RHE, 2 dpi with *S. meliloti*). The green MtAnn1-GFP fusion labels the nuclear periphery and cytoplasmic bridge (arrowhead) that forms in *ern1* root hairs at an early RHE stage. CFP-expressing *S. meliloti* bacteria (arrow) within the infection chamber are visualized in magenta. (**C**) Relative intensity of NR-GECO1 fluorescence, expressed as signal-to-noise ratio (SNR, cf. Methods section) in the RHE hair nucleus. (**D**-**E**) Confocal images illustrate MtAnn1-GFP green fluorescence labelling in the nuclear periphery and radiating cytoplasmic strands in outer cortical cells (C1a-c, C1f-g,

C1i) of an *ern1* infected root site (4 dpi with *S. meliloti*). (**E**) Maximal z-projection of the NR-GECO acquisition time series (see corresponding zone in **D**, grey dotted line). All outer cortical nuclei in focus are indicated (dashed white round/oval shapes). $Ca^{2+}$ spiking traces (**F**) reflect the intensity of NR-GECO1 fluorescence, expressed as signal-to-noise ratio (SNR, cf. Methods section) in cortical cells (**D**-**E**, C1a-C1j). Data for the *ern1* mutant are from 3 independent experiments (n=3). (**G**-**J**) Analysis of *dmi3* complemented with a *pEXT:DMI3* construct. (**G**-**H**) This complemented line exhibit restored root hair IT development (arrow), cytoplasmic bridge formation (arrowhead) and MtAnn1-GFP expression (green signal) (**G**). Adjacent outer cortical cells (CO) C1a to C1c (in **H**) do not show MtAnn-GFP labelling, in contrast to the A17 wid-type genotype (Fig. 3). The white asterisk indicates the position of the epidermal cell body of the infected root hair (in **G**-**H**, delimited by a dashed line in **G**). (**I**-**J**) Absence of cytoplasmic reorganization in outer cortical cells near rhizobia infected root hairs (**I**, arrowheads) in the complemented *dmi3* line is also evidenced by a constitutively expressed GFP transformation marker. White asterisks in **I**-**J** indicate the position of the epidermal cell bodies of infected root hairs. Complementation data (*dmi3* + *pEXT:DMI3*) were obtained at 6 dpi with CFP-expressing *S. meliloti* (magenta) from 3 independent experiments (n=11). Scale bars: **A**-**B** = 10 µm, **D**-**E**, **G**-**J** = 20 µm.

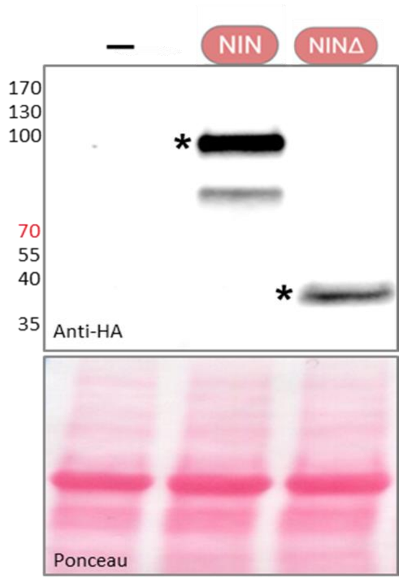


**Supplementary Fig. S10. Western blot shows the accumulation of 3x HA-tagged NIN versions in infiltrated *N. benthamiana* cells (related to Fig. 4).**

Transactivation studies using the *pMtAnn1:GUS* fusion were performed in *N. benthamiana* leaf discs with or without (-) co-infiltration with 3x HA-tagged NIN lacking its DNA binding domain (NINΔ) or not (NIN). The upper image shows Western blot analysis using anti-HA antibodies. NIN protein bands of expected sizes are indicated by asterisks. Molecular Weight sizes (in KDa) are indicated on the left. The bottom panel shows protein loading by Ponceau staining. See Fig. 4 for the corresponding transactivation experiments.

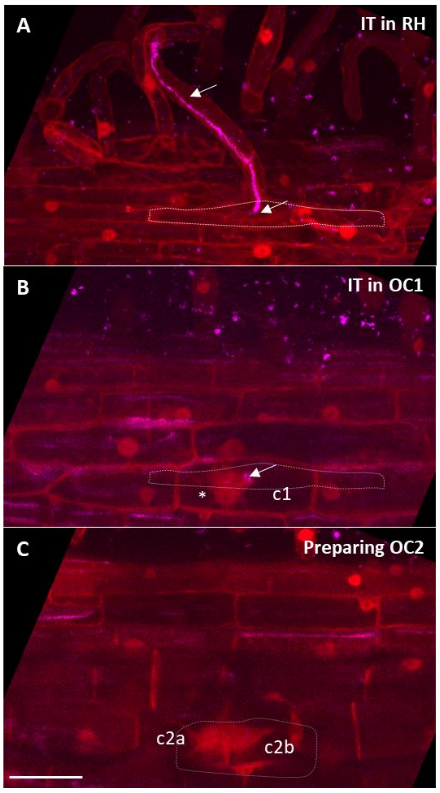


**Supplementary Fig. S11. Visualization of cytoplasmic bridge formation in *sunn* expressing a *pUBQ10*-driven DsRed marker (related to Fig. 4).**

Roots of *M. truncatula sunn* plants expressing *pUBQ10-DsRed* were inoculated with CFP-expressing *S. meliloti* (in magenta). The three images recapitulate the epidermal and two outer cortex cell layers at a rhizobial infection site. An infection thread reaching OC1 is shown (arrows). Images are merges of DsRed (in red) and CFP (in magenta). (**A**) shows the root hair infection thread (arrows) and the position of the epidermal body of the infected root hair (white dotted line). (**B**) corresponds to the adjacent OC1 cells, one of them (c1) hosting the infection thread (arrow) immersed in DsRed-labelled cytoplasm, close to the nucleus (asterisk). The position of the epidermal body of the infected root hair is represented by the grey dotted line. (**C**) focuses on the OC2 layer, in which two cells (c2a, c2b) directly adjacent to the infected OC1 cell (which contour is depicted by the grey dotted line) display cytoplasmic rearrangement highlighted by the DsRed fluorescence. Images in **A**-**C** were obtained from a single confocal stack, as maximal z-projections of sub-stacks comprising 11 (**A**), 11 (**B**) and 7 (**C**) successive sections, respectively. Scale bar = 40 µm.

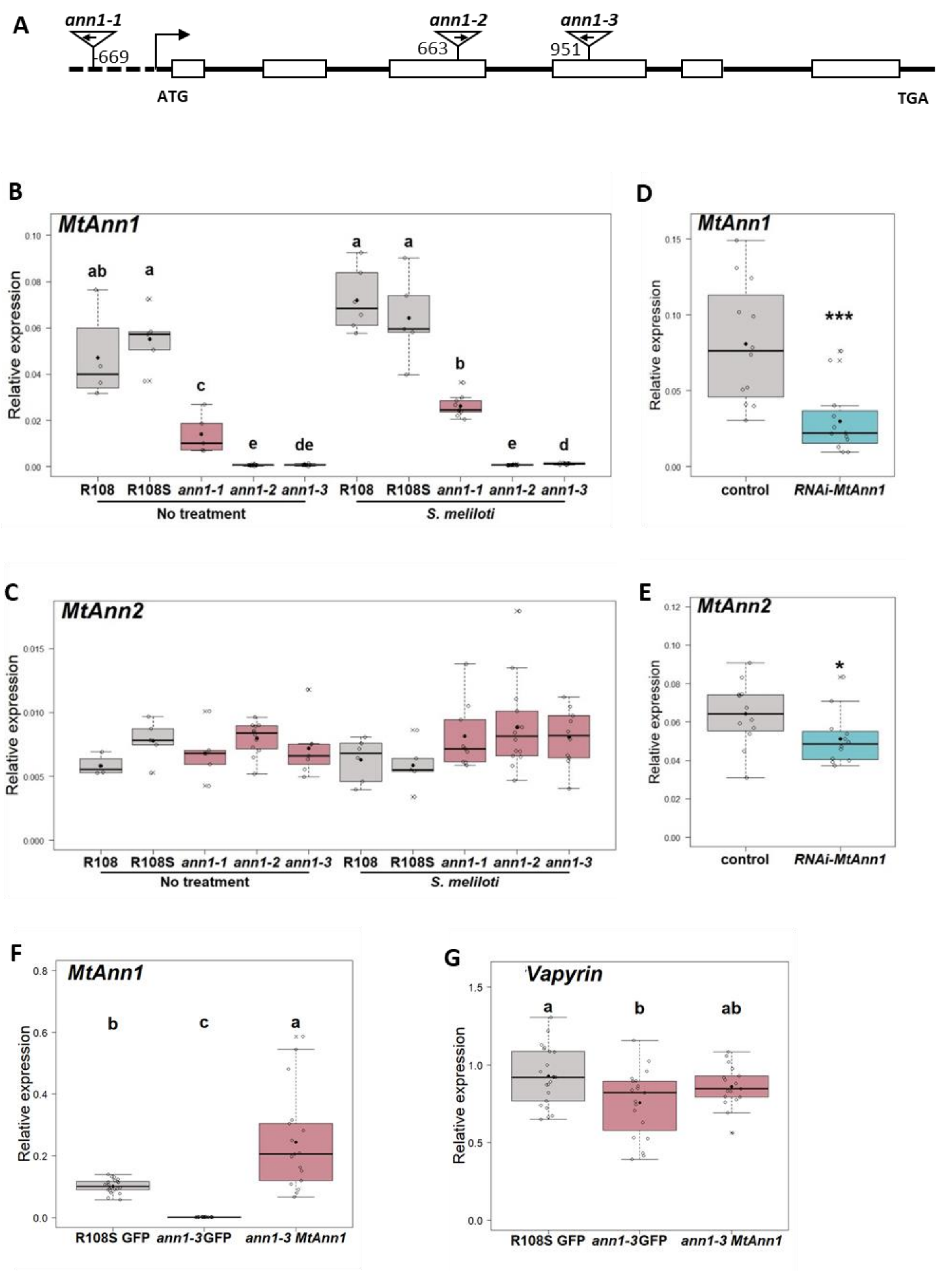


**Supplementary Fig. S12. Molecular characterization of *MtAnn1 Tnt1* insertion lines and RNAi roots (related to Figs. 5-6).**

(**A**) Schematic illustration of the *Tnt1* insertions in the *MtAnn1* gene. *ann1-1, ann1-2* and *ann1-3* insertions are respectively located -669 bp upstream the start codon, in exon 3 or 4. Exons (white rectangles) and introns (black lines) are represented at full scale, upstream regions or *Tnt1* inserts are not. (**B**-**G**) Q-RT-PCR expression analysis of *MtAnn1* (**B**, **D** and **F**), *MtAnn2* (**C** and **E**) and Vapyrin (**G**) in total RNA samples of non-inoculated (No treatment) or *S. meliloti* - inoculated roots (4 dpi, in **B**-**C** and **F**-**G**, and 5 dpi in **D**-**E**) from *ann1* mutant alleles (*ann1-3*, *ann1-2* and *ann1-3*) and WT control (R108 and/or R108S sibling) lines (**B**-**C**), from transgenic *RNAi-MtAnn1* and control (*pMtAnn1*-GUS) plants (**D**-**E**) and from transgenic WT R108S or *ann1-3* transformed with p35S-GFP (control) or p35S-MtAnn1-GFP (**F**-**G**). Expression values after normalization against the ubiquitin reference transcript are indicated. Box plots in **B**-**G** represent the distribution of individual values (open circles). Data in **B**-**C** is from 2 independent experiments n=6 (for R108), n=5 (for R108S), n=9 (for *ann1-1),* n=14 (for *ann1-2*) and n=10 (for *ann1-3*). Data in **D**-**E** is from 3 independent experiments n=12 (for control) and n=12 (for *RNAi-MtAnn1*), and data in **F**-**G** is from 2 independent experiments n=22 (for R108S p35S-GFP), n=19 (for *ann1-3* p35S-GFP) and n=17 (for *ann1-3* p35S-MtAnn1-GFP). Median (central line), mean (solid black circle) and outliers (crosses) are indicated. ANOVA followed by a Tukey HSD test in **B**-**C** and **G**, and a Kruskal Wallis test in **F** were performed in R. Classes sharing the same letter are not significantly different in ($p<0.0001$). Two-tailed t-tests were performed for values in **D**-**E** (* $p<0.05$, *** $p<0.001$).

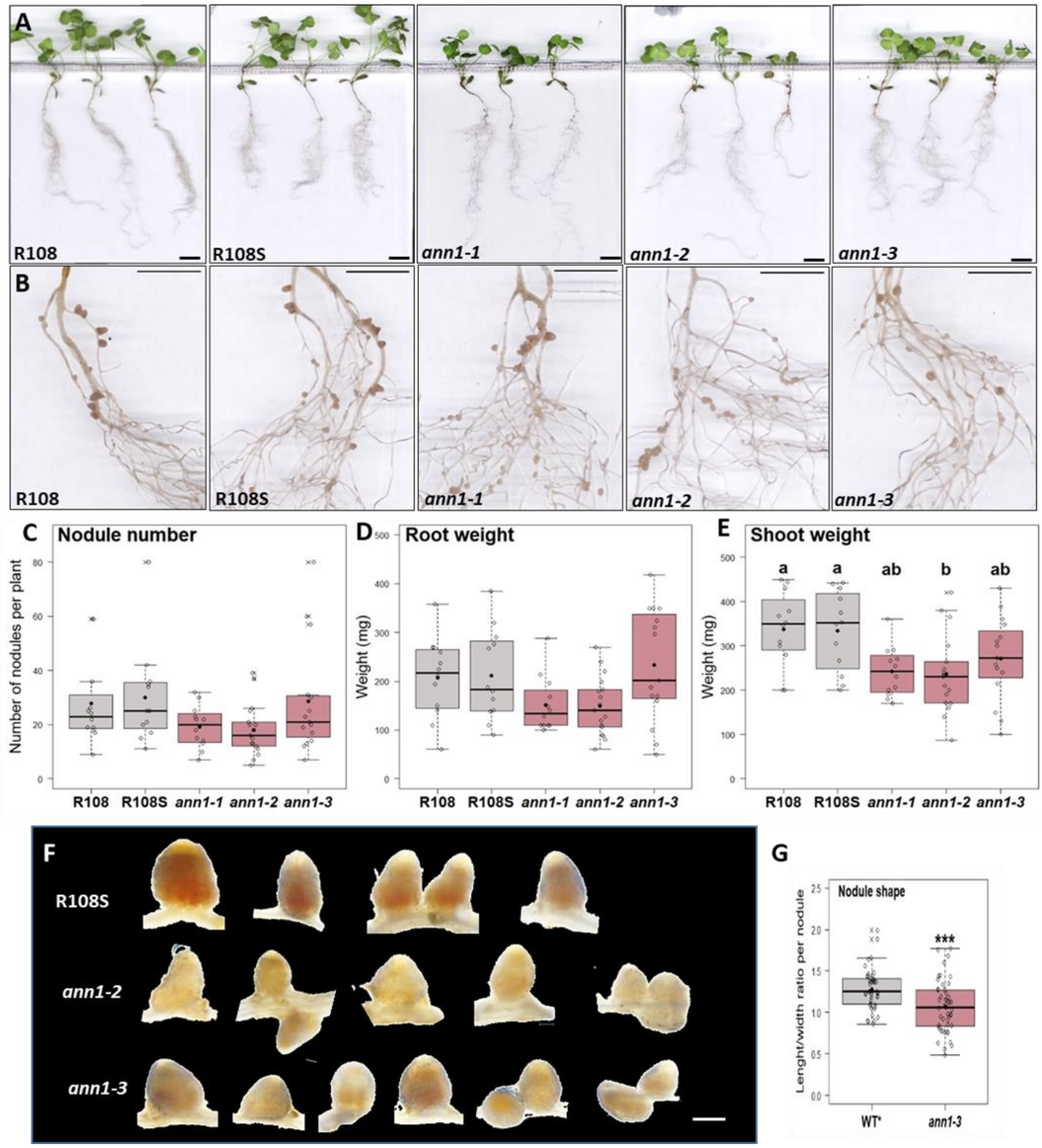


**Supplementary Fig. S13. Phenotypic characterization of *MtAnn1 Tnt1* insertion lines (related to Figs. 5-6).**

(**A**-**B** and **F**) Representative images of R108, R108S, *ann1-1*, *ann1-2* and *ann1-3* plants (**A**), their nodulated root systems (**B**) and nodules formed of these roots (**F**) and quantification of their shape (**G**) at 21 dpi. (**C**-**E**) Number of nodules per plant at 21 dpi (**C**), fresh root weight (**D**) and fresh shoot weight (**E**) in R108 (n=12), R108S (n=12), *ann1-1* (n=12), *ann1-2* (n=17) and *ann1-3* (n=15). (**G**) Nodule shape, represented by the ratio of maximal length to maximal width of individual nodules, was measured on R108S (n=43) and *ann1-3* (n=48) mutant nodules from 3 independent experiments. Box plots in **C**-**E** and **G** represent the distribution of individual values (open circles) from 3 independent experiments. Median (central line), mean (solid black circle) and outliers (crosses) are indicated. ANOVA (**C** and **E**), Kruskal Wallis (**D**) followed by Tukey HSD tests or a two-tailed t-test (**G**) were performed in R. Differences were not significant in **C**-**D** and asterisks in **G** indicate statistical difference (***p < 0.001). Classes sharing the same letter in **E** are not significantly different (α=5%). Scale bars: **A** = 2 cm, **B** = 1 cm, **F** = 2.5 mm.

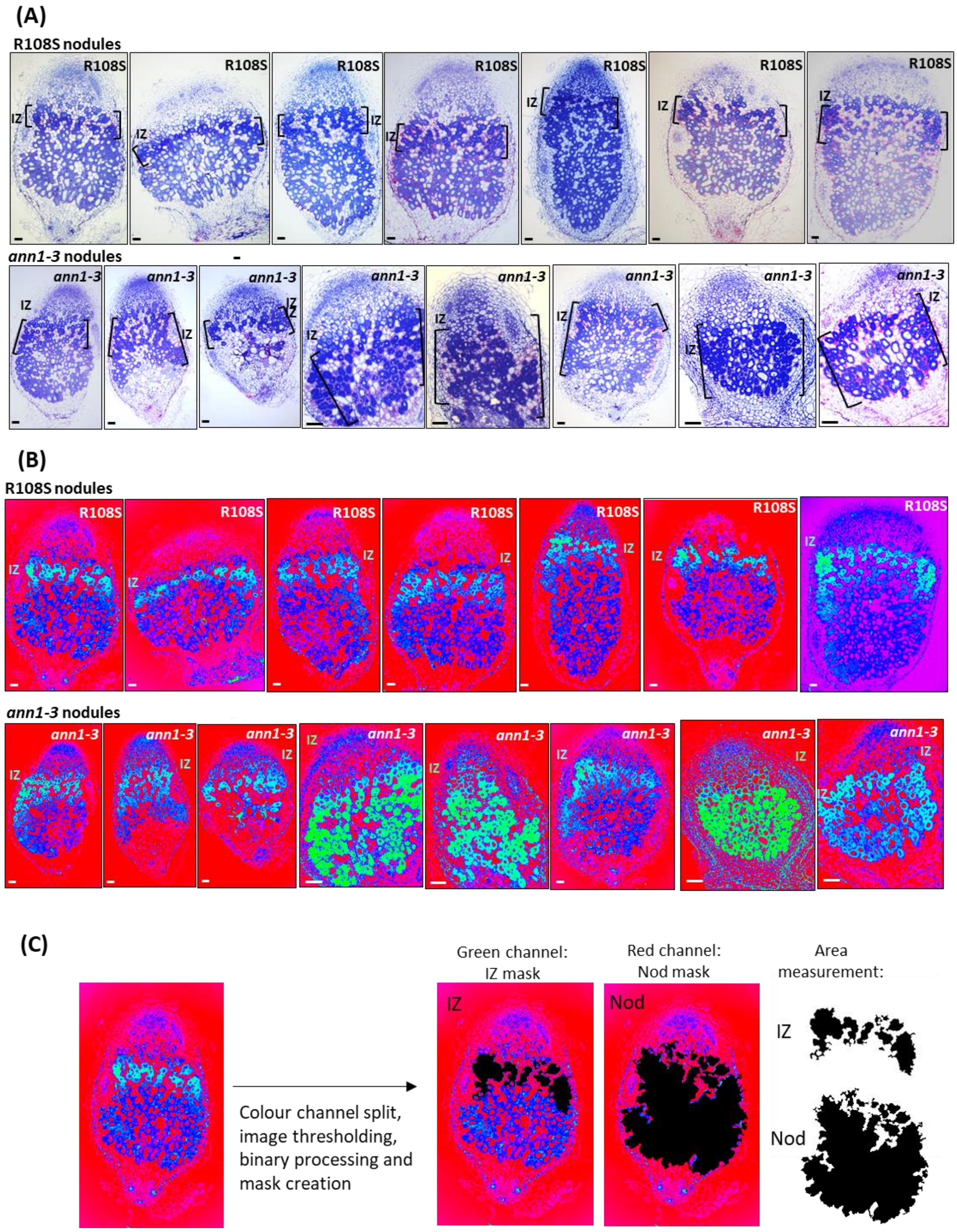


**Supplementary Fig. S14. Comparative analysis of nodule zones in R108 and *ann1-3* nodules by ImageJ (related to Fig. 6).**

(**A**-**B**) To measure the relative area of nodule interzones (IZ), bright-field images (**A**) of toluidine blue/basic fuchsin-stained 1 µm longitudinal sections of nodules from R108S and *ann1-3* mutant roots at 21 dpi with *S. meliloti* were first transformed into spectrum-type image (**B**) in ImageJ, which allows better visualization of nodule IZs (in light blue to light green according to signal intensity). (**C**) Schematic representation of the ImageJ procedure used to measure the area of respective IZs and remaining nodule zones (Nod). First, spectrum images were color split and the resulting green channel (with IZ zone) and red channel (with proximal ZII, IZ and ZIII of nodules, called Nod) images were selected for further processing. Image thresholding and binary processing (fill holes) were applied to selected images to generate corresponding IZ and Nod masks (shown in black) that were then used for area measurement in ImageJ. The resulting data for IZ/Nod area per nodule is shown in Fig. 6P.

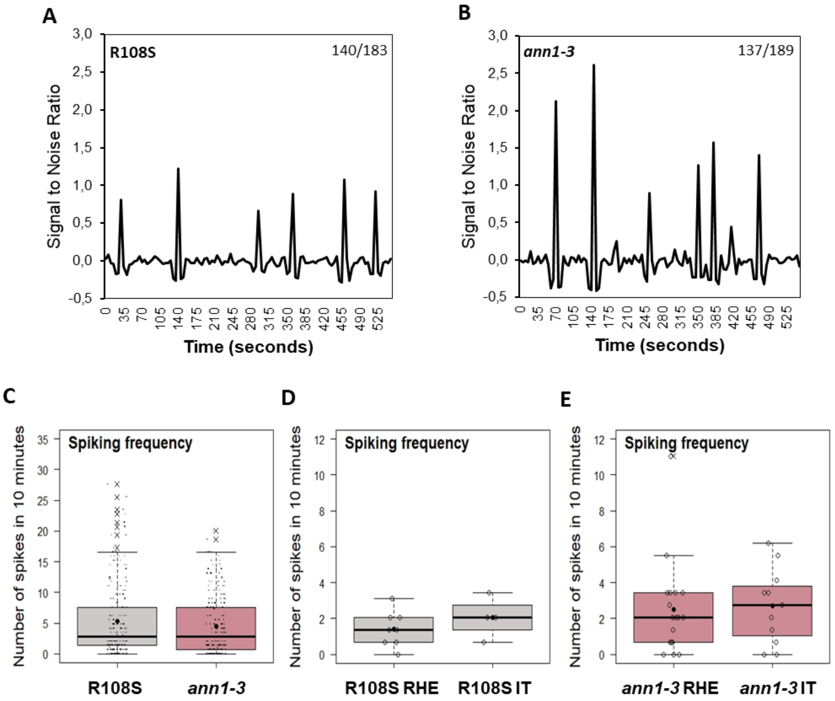


**Supplementary Fig. S15. Nuclear $Ca^{2+}$ spiking patterns in the absence of *MtAnn1* (related to Fig. 7).**

Transgenic roots expressing the NR-GECO1 $Ca^{2+}$ sensor were generated to monitor nuclear $Ca^{2+}$ oscillations in R108S (wild-type) and in the *ann1-3* mutant. (**A**-**B**) Representative nuclear $Ca^{2+}$ spiking traces of R108S (**A**) or *ann1-3* (**B**) in non-infected root hairs inoculated with *S. meliloti* at 1-4 dpi. Variations in red fluorescence of the NR-GECO1 $Ca^{2+}$ sensor are reported as the Signal to Noise Ratio (SNR, cf. Methods section). Values on the top right correspond to the number of root hairs showing more than 2 $Ca^{2+}$ spikes in 10 minutes, out of total root hairs analyzed. (**C**-**E**) Spiking frequencies represent the number of spikes in 10 minutes for each nucleus in *S. meliloti*-responsive, non-infected root hairs (R108S, n=183 and *ann1-3*, n=189) (**C**) or in those containing entrapped rhizobia (RHE) or ITs in R108S (R108S RHE, n=8; R108S with ITs=4) (**D**) and *ann1-3* (*ann1-3* RHE=18; *ann1-3* root hairs with ITs= 11) (**E**) backgrounds. Two-tailed Mann-Whitney tests of values did not reveal statistical differences ($\alpha$=5%).

**Supplementary Fig. S16. Conservation of *MtAnn1* in root endosymbioses (related to Fig. 8).**

Protein sequence of MtAnn1 (MtrunA17_Chr8g0352611) was used as query to search against a database containing 227 plant genomes covering the main lineages of green plants and five SAR (*Stramenopiles*/*Alveolata*/*Rhizaria*) genomes as outgroups[57]. Homologous proteins were then aligned using Muscle v5.1 and the phylogenetic tree was constructed using the NJ method. Plant species establishing (filled triangles, circles or squares) or not establishing (empty triangles, circles or squares) endosymbioses (orange squares), AM symbiosis (blue circles) or rhizobia root nodule symbiosis (RNS) (orange triangles) are indicated.

**Table S1. Primers used in this study.**

**ann1 mutants genotyping primers**

| | |
|---|---|
| ***LTR4-*** | TACCGTATCTCGGTGCTACA |
| ***LTR6-*** | GCTACCAACCAAACCAAGTCAA |
| ***Mtr14183-Rev*** | CTCTGCTCACAATCACACGG |
| ***-344 pMtAnn1-Fw*** | TGATGATTTGAGTCTGGTCCCA |
| ***-914 pMtAnn1-Fw*** | ACCGTCCCTGTGACATTGAT |

**GoldenGate cloning primers**

| | |
|---|---|
| ***pMtAnn1-GG-A-Fw*** | **GGTCTC**GAAATGAATTCTTTTATTGATTGCG |
| ***pMtAnn1-GG-B-Rev*** | **GGTCTC**GTTTGgGTTCAATATGTaTTGTTTATAg |
| ***MtAnn1-ATG-GG-B-Fw*** | **GGTCTC**TCAAAATGGCTACCCTTTCTGCTCCTAG |
| ***MtAnn1-STOP-GG-C-Rev*** | **GGTCTC**GCACCTCATTCCTCTTTCCCCAAGAGAG |
| ***MtAnn1 antisens-STOP-GG-X-Fw*** | **GGTCTC**GCCAGTCATTCCTCTTTCCCCAAGAGAG |
| ***MtAnn1 antisens-ATG-GG-D-Rev*** | **GGTCTC**GCGTAATGGCTACCCTTTCTGCTCCTAG |
| ***GUS-GG-B-Fw*** | **GGTCTC**TCAAAATGGTCCGTCCTGTAGAAACC |
| ***GUS-GG-D-Rev*** | **GGTCTC**ACGTATCATTGTTTGCCTCCCTGCTG |

**qRT-PCR primers**

| | |
|---|---|
| ***1271 Ubiquitin-Fw*** | TTGTGTGTTGAATCCTAAGCAGG |
| ***1334 Ubiquitin-Rev*** | CAAGACCCATGCAACAAGTTCT |
| ***2247 Chr6g0453261-Fw*** | GTTCTTGTTGCTGCCAAATACC |
| ***2359 Chr6g0453261-Rev*** | CTGCTCCGTTTCATAAACCTTG |
| ***-5 MtAnn1-Fw*** | GAACTATGGCTACCCTTTCTGC |
| ***88 MtAnn1-Rev*** | CATCAGTACCCCATCCTTCG |

**Table S2. List of plant genomes used for the phylogeny analysis.**

| **species_code** | **species_name** | **lineage** | **species_order** | **species_family** | **source** | **endosymbiosis** | **AMS** | **RNS** |
|---|---|---|---|---|---|---|---|---|
| Abrpre | Abrus precatorius | Angiosperms | Fabales | Fabaceae | NCBI | 1 | 1 | 1 |
| Adicap | Adiantum capillus-veneris | Monilophytes | Polypodiales | Polypodiaceae | 10.1038/s41477-022-01222-x | 1 | 1 | 0 |
| Adinel | Adiantum | Monilop | Polypod | Polypodia | 10.1038/gbe/ev | 1 | 1 | 0 |
| Aeseve | Aeschynomene evenia | Angiosperms | Fabales | Fabaceae | 10.1038/s41467-021-21094-7 | 1 | 1 | 1 |
| Alnglu | Alnus glutinosa | Angiosperms | Fagales | Betulaceae | 10.1126/science.aat1743 | 1 | 1 | 1 |
| Alsspi | Alsophila spinulosa | Monilophytes | Cyatheales | Cyatheaceae | 10.1038/s41477-022-01146-6 | 1 | 1 | 0 |
| Amahyp | Amaranthus hypochondriacus | Angiosperms | Caryophyllales | Amaranthaceae | 10.3835/plantgenome2015.07.0062 | 0 | 0 | 0 |
| Ambtri | Amborella trichopoda | Angiosperms | Amborellaceae | Amborellales | 10.1126/science.1241089 | 1 | 1 | 0 |
| Anacom | Ananas comosus | Angiosperms | Poales | Bromeliaceae | 10.1038/ng.3435 | 1 | 1 | 0 |
| Antagr BONN | Anthoceros agrestis cv. BONN | Hornworts | Anthocerotales | Anthocerataceae | 10.1038/s41477-020-0618-2 | 1 | 1 | 0 |
| Antagr OXF | Anthoceros agrestis cv. OXF | Hornworts | Anthocerotales | Anthocerataceae | 10.1038/s41477-020-0618-2 | 1 | 1 | 0 |
| Antpun | Anthoceros punctatus | Hornworts | Anthocerotales | Anthocerataceae | 10.1038/s41477-020-0618-2 | 1 | 1 | 0 |
| Apafus SAG2523 | Apatococcus fuscideae SAG2523 | Chlorophytes | Chlorellales | Chlorellaceae | 10.1101/2022.01.06.475074 | 0 | 0 | 0 |
| Apalob SAG2145 | Apatococcus lobatus | Chlorophytes | Chlorellales | Chlorellaceae | Phytozome | 0 | 0 | 0 |
| Aposhe | Apostasia shenzhenica | Angiosperms | Asparagales | Orchidaceae | 10.1038/nature23897 | 1 | 0 | 0 |
| Aqucoe | Aquilegia coerulea | Angiosperms | Ranunculales | Ranunculaceae | 10.7554/eLife.36426 | 1 | 1 | 0 |
| Aradur | Arachis duranensis | Angiosperms | Fabales | Fabaceae | 10.1038/ng.3517 | 1 | 1 | 1 |
| Arahal | Arabidopsis halleri | Angiosperms | Brassicales | Brassicaceae | 10.5061/dryad.gn4hh | 0 | 0 | 0 |

| | | | | | | | | |
|---|---|---|---|---|---|---|---|---|
| Arahyp | Arachis hypogaea | Angiosperms | Fabales | Fabaceae | 10.25739/hb5x-wx74 | 1 | 1 | 1 |
| Araipa | Arachis ipaensis | Angiosperms | Fabales | Fabaceae | 10.1038/ng.3517 | 1 | 1 | 1 |
| Aralyr | Arabidopsis lyrata | Angiosperms | Brassicales | Brassicaceae | 10.1038/ng.807 | 0 | 0 | 0 |
| Aratha | Arabidopsis thaliana | Angiosperms | Brassicales | Brassicaceae | 10.1093/nar/gkr1090 | 0 | 0 | 0 |
| Astglo | Asterochloris glomerata | Chlorophytes | Trebouxiales | Trebouxiaceae | 10.1186/s12864-019-5629-x | 0 | 0 | 0 |
| Auxpro | Auxenochlorella protothecoides | Chlorophytes | Chlorellales | Chlorellaceae | 10.1186/1471-2164-15-582 | 0 | 0 | 0 |
| Auxpro UTEX25 | Auxenochlorella protothecoides UTEX25 | Chlorophytes | Chlorellales | Chlorellaceae | NCBI | 0 | 0 | 0 |
| Azofil | Azolla filiculoides | Monilophytes | Salviniales | Salviniaceae | 10.1038/s41477-018-0188-8 | 0 | 0 | 0 |
| Batpra | Bathycoccus prasinos RCC1105 | Chlorophytes | Mamiellales | Bathycoccaceae | 10.186/gb-2012-13-8-r74 & 10.1371/journal.pone.0039648 | 0 | 0 | 0 |
| Bauvar | Bauhinia variegata | Angiosperms | Fabales | Fabaceae | 10.1093/dnares/dsac012 | 1 | 1 | 0 |
| Begfuc | Begonia fuchsioides | Angiosperms | Cucurbitales | Begoniaceae | 10.1126/science.aat1743 | 1 | 1 | 0 |
| Benhis | Benincasa hispida | Angiosperms | Cucurbitales | Cucurbitaceae | 10.1038/s41467-019-13185-3 | 1 | 1 | 0 |
| Betpat | Beta patula | Angiosperms | Caryophyllales | Amaranthaceae | 10.1111/tpj.14413 | 0 | 0 | 0 |
| Betpen | Betula pendula | Angiosperms | Fagales | Betulaceae | 10.1038/ng.3862 | 1 | 1 | 0 |
| Betvul | Beta vulgaris | Angiosperms | Caryophyllales | Amaranthaceae | 10.1111/2020.09.15.298315 | 0 | 0 | 0 |
| Boestr | Boechera stricta | Angiosperms | Brassicales | Brassicaceae | Phytozome | 0 | 0 | 0 |
| Botbra | Botryococcus braunii | Chlorophytes | Chlorellales | Botryococcaceae | NA | 0 | 0 | 0 |
| Bradis | Brachypodium distachyon | Angiosperms | Poales | Poaceae | 10.1038/nature08747 | 1 | 1 | 0 |
| Braolecap | Brassica oleraceae capitata | Angiosperms | Brassicales | Brassicaceae | 10.1038/ncomms4930 | 0 | 0 | 0 |

| | | | | | | | | |
|---|---|---|---|---|---|---|---|---|
| Brarap | Brassica rapa FPsc | Angiosperms | Brassicales | Brassicaceae | 10.1038/s41438-018-0071-9 | 0 | 0 | 0 |
| Cajcaj | Cajanus cajan | Angiosperms | Fabales | Fabaceae | 10.1038/nbt.2022. | 1 | 1 | 1 |
| Camsin | Camellia sinensis | Angiosperms | Ericales | Theaceae | NCBI | 1 | 1 | 0 |
| Cansat | Cannabis sativa | Angiosperms | Rosales | Cannabaceae | NCBI | 1 | 1 | 0 |
| Capann | Capsicum annuum | Angiosperms | Solanales | Solanaceae | 10.1073/pnas.1400975111 | 1 | 1 | 0 |
| Capgra | Capsella grandiflora | Angiosperms | Brassicales | Brassicaceae | 10.1038/ng.2669 | 0 | 0 | 0 |
| Caprub | Capsella rubella | Angiosperms | Brassicales | Brassicaceae | 10.1038/ng.2669 | 0 | 0 | 0 |
| Carfan | Carpinus fangiana | Angiosperms | Fagales | Betulaceae | 10.1038/s41597-020-0370-5 | 1 | 1 | 0 |
| Carlit | Carex littledalei | Angiosperms | Poales | Cyperaceae | NCBI | 0 | 0 | 0 |
| Carpap | Carica papaya | Angiosperms | Brassicales | Caricaceae | 10.1038/nature06856 | 1 | 1 | 0 |
| Casaus | Castanospermum australe | Angiosperms | Fabales | Fabaceae | 10.1126/science.aat1743 | 1 | 1 | 0 |
| Casgla | Casuarina glauca | Angiosperms | Fagales | Casuarinaceae | 10.1111/tpj.16201 | 1 | 1 | 1 |
| Casmol | Castanea mollissima | Angiosperms | Fagales | Fagaceae | NCBI | 1 | 1 | 0 |
| Cepfol | Cephalotus follicularis | Angiosperms | Oxalidales | Cephalotaceae | 10.1038/s41559-016-0059 | 0 | 0 | 0 |
| Cercan | Cercis canadensis | Angiosperms | Fabales | Fabaceae | 10.1126/science.aat1743 | 1 | 1 | 0 |
| Cerpur | Ceratodon purpureus | Mosses | Dicranales | Dicranaceae | NCBI | 0 | 0 | 0 |
| Cerric | Ceratopteris richardii | Monilophytes | Polypodiales | Pteridaceae | 10.1038/s41477-022-01226-7 | 0 | 0 | 0 |
| Chabra | Chara braunii | Charophytes | Charales | Characeae | 10.1016/j.cell.2018.06.033 | 0 | 0 | 0 |
| Chafas | Chamaecrista fasciculata | Angiosperms | Fabales | Fabaceae | 10.1126/science.aat1743 | 1 | 1 | 1 |
| Chequi | Chenopodium quinoa | Angiosperms | Caryophyllales | Chenopodiaceae | 10.1038/nature21370 | 0 | 0 | 0 |
| Chlatm | Chlorokybus atmophyticus | Charophytes | Chlorokybales | Chlorokybaceae | 10.1038/s41477-019-0560-3 | 0 | 0 | 0 |
| Chlrei | Chlamydomonas reinhardtii | Chlorophytes | Chlamydomonales | Chlamydomonaceae | 10.1126/science.1143609 | 0 | 0 | 0 |

| | | | | | | | | |
|---|---|---|---|---|---|---|---|---|
| Chlsor1602 | Chlorella sorokiniana 1602 | Chlorophytes | Chlorellales | Chlorellaceae | NCBI | 0 | 0 | 0 |
| Chlvar | Chlorella variabilis | Chlorophytes | Chlorellales | Chlorellaceae | 10.1105/tpc.110.076406 | 0 | 0 | 0 |
| Chocri | Chondrus crispus | SAR | Gigartinales | Gigartinaceae | 10.1073/pnas.1221259110 | 0 | 0 | 0 |
| Chrzof | Chromochloris zofingiensis | Chlorophytes | Sphaeropleales | Chromochloridaceae | 10.1073/pnas.1619928114 | 0 | 0 | 0 |
| Cicari | Cicer arietinum ICC4958 | Angiosperms | Fabales | Fabaceae | 10.1038/SREP12806 | 1 | 1 | 1 |
| Citcle | Citrus clementina | Angiosperms | Sapindales | Rutaceae | 10.1038/nbt.2906 | 1 | 1 | 0 |
| Citlan | Citrullus lanatus ssp vulgaris 97103 | Angiosperms | Cucurbitales | Cucurbitaceae | 10.1038/ng.2470 | 1 | 1 | 0 |
| Citsin | Citrus sinensis | Angiosperms | Sapindales | Rutaceae | 10.1038/nbt.2906 | 1 | 1 | 0 |
| CocpriSAG2167 | Coccomyxa pringsheimii SAG216-7 | Chlorophytes | Incertae sedis | Coccomyxaceae | 10.1101/2022.01.06.475074 | 0 | 0 | 0 |
| CocsubC169 | Coccomyxa subellipsoidea C-169 | Chlorophytes | Incertae sedis | Coccomyxaceae | 10.1186/gb-2012-13-5-r39 | 0 | 0 | 0 |
| Cucargarg | Cucurbita argyrosperma ssp argyrosperma | Angiosperms | Cucurbitales | Cucurbitaceae | NCBI | 1 | 1 | 0 |
| Cucmax | Cucurbita maxima | Angiosperms | Cucurbitales | Cucurbitaceae | 10.1016/j.molp.2017.09.003 | 1 | 1 | 0 |
| Cucmel | Cucumis melo | Angiosperms | Cucurbitales | Cucurbitaceae | 10.1073/pnas.1205415109 | 1 | 1 | 0 |
| Cucmos | Cucurbita moschata | Angiosperms | Cucurbitales | Cucurbitaceae | 10.1016/j.molp.2017.09.003 | 1 | 1 | 0 |
| Cucpep | Cucurbita pepo | Angiosperms | Cucurbitales | Cucurbitaceae | 10.1111/pbi.12860 | 1 | 1 | 0 |
| Cucsat | Cucumis sativus PI183967 | Angiosperms | Cucurbitales | Cucurbitaceae | 10.1016/j.molp.2017.09.003 | 1 | 1 | 0 |
| Cuscam | Cuscuta campestris | Angiosperms | Solanales | Convolvulaceae | NCBI | 0 | 0 | 0 |
| Cyapar | Cyanophora paradoxa | Glaucophytes | Glaucocustales | Glaucocystaceae | 10.1126/science.1213561 | 0 | 0 | 0 |

| | | | | | | | | |
|---|---|---|---|---|---|---|---|---|
| Cycmic | Cycas micholitzii | Gymnosperms | Cycadales | Cycadaceae | Gymno-plaza | 1 | 1 | 0 |
| Datglo | Datisca glomerata | Angiosperms | Cucurbitales | Datiscaceae | 10.1126/science.aat1743 | 1 | 1 | 1 |
| Daucar | Daucus carota | Angiosperms | Apiales | Apiaceae | 10.1038/ng.3565 | 1 | 1 | 0 |
| Dencat | Dendrobium catenatum | Angiosperms | Asparagales | Orchidaceae | 10.1038/nature23897 | 1 | 0 | 0 |
| Diacar | Dianthus caryophyllus | Angiosperms | Caryophyllales | Caryophyllaceae | 10.1093/dnares/dst053 | 0 | 0 | 0 |
| Distri | Discaria trinervis | Angiosperms | Rosales | Rhamnaceae | 10.1126/science.aat1743 | 1 | 1 | 1 |
| Drydru | Dryas drummondii | Angiosperms | Rosales | Rosaceae | 10.1126/science.aat1743 | 1 | 1 | 1 |
| Dunsal | Dunaliella salina | Chlorophytes | Chlamydomonales | Dunaliellaceae | 10.1128/genomeA.01105-17 | 0 | 0 | 0 |
| EllbilSAG24580 | Elliptochloris bilobata SAG245-80 | Chlorophytes | Prasiolales | incertae sedis | 10.1101/2022.01.06475074 | 0 | 0 | 0 |
| Entsed | Entodon seductrix | Mosses | Hypnales | Entodontaceae | 10.1093/gbe/evac020 | 0 | 0 | 0 |
| Eurfer | Euryale ferox | Angiosperms | Nymphaeales | Nymphaeaceae | NCBI | 0 | 0 | 0 |
| Eutsal | Eutrema salsugineum | Angiosperms | Brassicales | Brassicaceae | 10.3389/fpls.2013.00046 | 0 | 0 | 0 |
| Faialb | Faidherbia albida | Angiosperms | Fabales | Fabaceae | 10.1093/gigascience/giy152 | 1 | 1 | 1 |
| Fonant | Fontinalis antipyretica | Mosses | Hypnales | Fontinalaceae | 10.5524/100748 | 0 | 0 | 0 |
| Fraana | Fragaria x ananassa Camarosa | Angiosperms | Rosales | Rosaceae | 10.1038/s41438-0.21-00476-4 | 1 | 1 | 0 |
| Fraexc | Fraxinus excelsior | Angiosperms | Lamiales | Oleaceae | 10.1038/nature20786 | 1 | 1 | 0 |
| Fraiin | Fragaria iinumae | Angiosperms | Rosales | Rosaceae | NCBI | 1 | 1 | 0 |
| Fraves | Fragaria vesca | Angiosperms | Rosales | Rosaceae | 10.1093/gigascience/gix124 | 1 | 1 | 0 |
| Galsul | Galdieria sulphuraria | SAR | Cyanidiales | Galdieriaceae | 10.1126/science.1231707 | 0 | 0 | 0 |
| Ginbil | Ginkgo biloba | Gymnosperms | Ginkgoales | Ginkgoceae | 10.5524/100613 | 1 | 1 | 0 |
| Glymax | Glycine max | Angiosperms | Fabales | Fabaceae | 10.1038/nature08670 | 1 | 1 | 1 |
| Glysoj | Glycine soja | Angiosperms | Fabales | Fabaceae | NCBI | 1 | 1 | 1 |

| | | | | | | | | |
|---|---|---|---|---|---|---|---|---|
| Gnemon | Gnetum montanum | Gymnosperms | Gnetales | Gnetaceae | 10.5061/dryad.0vm37 | 1 | 1 | 0 |
| Gosrai | Gossypium raimondii | Angiosperms | Malvales | Malvaceae | 10.1038/nature11798 | 1 | 1 | 0 |
| Helann | Helianthus annuus | Angiosperms | Asterales | Asteraceae | 10.1038/nature22380 | 1 | 1 | 0 |
| HelspATCC50920 | Helicosporidium sp. ATCC 50920 | Chlorophytes | Chlorellales | Chlorellaceae | NCBI | 0 | 0 | 0 |
| Hevbra | Hevea brasiliensis | Angiosperms | Malpighiales | Euphorbiaceae | NCBI | 1 | 1 | 0 |
| Humlup | Humulus lupulus | Angiosperms | Rosales | Cannabaceae | 10.1093/pcp/pcu169 | 1 | 1 | 0 |
| Hypcur | Hypnum curvifolium | Mosses | Hypnales | Hypnaceae | 10.1093/gbe/evac020 | 0 | 0 | 0 |
| Isotai | Isoetes taiwanensis | Lycophytes | Isoetales | Isoetaceae | 10.1038/s41467-021-26644-7 | 1 | 1 | 0 |
| Jatcur | Jatropha curcas | Angiosperms | Malpighiales | Euphorbiaceae | NCBI | 1 | 1 | 0 |
| Jugreg | Juglans regia | Angiosperms | Fagales | Juglandaceae | 10.1111/tpj.13207 | 1 | 1 | 0 |
| Klenit | Klebsormidium nitens | Charophytes | Klebsormidiales | Klebsormidiaceae | 10.1038/ncomms4978 | 0 | 0 | 0 |
| Labpur | Lablab purpureus | Angiosperms | Fabales | Fabaceae | 10.1093/gigascience/giy152 | 1 | 1 | 1 |
| Lagsic | Lagenaria siceraria | Angiosperms | Cucurbitales | Cucurbitaceae | 10.1111/tpj.13722 | 1 | 1 | 0 |
| Linusi | Linum usitatissimum | Angiosperms | Malpighiales | Linaceae | 10.1111/j.1365-313X.2012.05093.x | 1 | 1 | 0 |
| Lotjap | Lotus japonicus | Angiosperms | Fabales | Fabaceae | 10.1101//2020.05.29.124313 | 1 | 1 | 1 |
| LuncruJGI | Lunularia cruciata | Liverworts | Lunulariales | Lunulariaceae | JGI | 1 | 1 | 0 |
| Lupalb | Lupinus albus | Angiosperms | Fabales | Fabaceae | 10.1038/s41467-019-14197-9 | 1 | 0 | 1 |
| Lupang | Lupinus angustifolius | Angiosperms | Fabales | Fabaceae | 10.1111/pbi.12615 | 1 | 0 | 1 |
| Lyccla | Lycopodium clavatum | Lycophytes | Lycopodiales | Lycopodiaceae | 10.1101/2022.12.06.519249 | 0 | 0 | 0 |
| Malbac | Malus baccata | Angiosperms | Rosales | Rosaceae | NCBI | 1 | 1 | 0 |
| Maldom | Malus domestica GDDH13_1_1 | Angiosperms | Rosales | Rosaceae | 10.1038/ng.3886 | 1 | 1 | 0 |

| | | | | | | | | |
|---|---|---|---|---|---|---|---|---|
| Manesc | Manihot esculenta | Angiosperms | Malpighiales | Euphorbiaceae | 10.1038/nbt.3535 | 1 | 1 | 0 |
| Marinf | Marchantia inflexa | Liverworts | Marchantiales | Marchantiaceae | 10.1038/s41598-019-45039-9 | 1 | 1 | 0 |
| Marpal | Marchantia paleaceae | Liverworts | Marchantiales | Marchantiaceae | 10.1126/science.abg0929 | 1 | 1 | 0 |
| Marpolmon | Marchantia polymorpha ssp. montivagans SA2 | Liverworts | Marchantiales | Marchantiaceae | 10.3389/fpls.2020.00829 | 0 | 0 | 0 |
| Marpolpol | Marchantia polymorpha ssp. polymorpha BR5 | Liverworts | Marchantiales | Marchantiaceae | 10.3389/fpls.2020.00829 | 0 | 0 | 0 |
| Marpolrud | Marchantia polymorpha ssp. ruderalis TAK1 v6.1 | Liverworts | Marchantiales | Marchantiaceae | 10.1016/j.cub.2019.12.015 | 0 | 0 | 0 |
| Medtru | Medicago truncatula | Angiosperms | Fabales | Fabaceae | 10.1038/s41477-018-0286-7 | 1 | 1 | 1 |
| Mesend | Mesotaenium endlicherianum | Charophytes | Zygnemales | Zygnematophyceae | 10.1016/j.cell.2019.10.019 | 0 | 0 | 0 |
| Mesvir | Mesostigma viride | Charophytes | Mesostigmatales | Mesostigmataceae | 10.1038/s41477-019-0560-3 | 0 | 0 | 0 |
| Miccon24180 | Micratinium conductrix SAG241.80 | Chlorophytes | Chlorellales | Chlorellaceae | NCBI | 0 | 0 | 0 |
| Micpus1545 | Micromonas pusilla CCMP1545 | Chlorophytes | Mamiellales | Mamiellaceae | 10.126/science.1167222 | 0 | 0 | 0 |
| MicpusNOUM17 | Micromonas pusilla NOUM17 | Chlorophytes | Mamiellales | Mamiellaceae | 10.126/science.1167222 | 0 | 0 | 0 |
| Mimpud | Mimosa pudica | Angiosperms | Fabales | Fabaceae | Intern | 1 | 1 | 1 |
| Momcha | Momordica charantia | Angiosperms | Cucurbitales | Cucurbitaceae | NCBI | 1 | 1 | 0 |
| Monneg | Monoraphidium neglectum | Chlorophytes | Sphaeropleales | Selenastraceae | 10.1186/1471-2164-14-926 | 0 | 0 | 0 |

| | | | | | | | | |
|---|---|---|---|---|---|---|---|---|
| Mornot | Morus notabilis | Angiosperms | Rosales | Moraceae | 10.1038/ncomms3445 | 1 | 1 | 0 |
| Morole | Moringa oleifera | Angiosperms | Brassicales | Moringaceae | 10.1093/gigascience/giy152 | 1 | 1 | 0 |
| Mucpru | Mucuna pruriens | Angiosperms | Fabales | Fabaceae | NCBI | 1 | 1 | 1 |
| Musacu | Musa acuminata | Angiosperms | Zingiberales | Zingiberaceae | 10.1093/database/bat035 | 1 | 1 | 0 |
| Myrbis SAG2043 | Myrmecia bisecta SAG2043 | Chlorophytes | Trebouxiales | Trebouxiaceae | 10.1101/2022.01.06.475074 | 0 | 0 | 0 |
| Nelnuc | Nelumbo nucifera | Angiosperms | Proteales | Nelumbonaceae | 10.1186/gb-2013-14-5-r41 | 0 | 0 | 0 |
| Nicben | Nicotiana benthamiana | Angiosperms | Solanales | Solanaceae | 10.1094/MPMI-06-12-0148-TA | 1 | 1 | 0 |
| Nissch | Nissolia schottii | Angiosperms | Fabales | Fabaceae | 10.1126/science.aat1743 | 1 | 1 | 0 |
| Nymcol | Nymphea colarata | Angiosperms | Nymphaeales | Nymphaeaceae | NCBI | 0 | 0 | 0 |
| Orysat | Oryza sativa | Angiosperms | Poales | Poaceae | 10.1093/nar/gkl976 | 1 | 1 | 0 |
| Ostluc | Ostreococcus lucimarinus | Chlorophytes | Mamiellales | Mamiellaceae | 10.1073/pnas.0611046104 | 0 | 0 | 0 |
| Osttau | Ostreococcus tauri | Chlorophytes | Mamiellales | Mamiellaceae | 10.1186/1471-2164-15-1103 & 10.1073/pnas.0611046104 | 0 | 0 | 0 |
| Parand | Parasponia andersonii | Angiosperms | Rosales | Cannabaceae | 10.1073/pnas.1721395115 | 1 | 1 | 1 |
| Parkes | Parachlorella kessleri | Chlorophytes | Chlorellales | Chlorellaceae | NCBI | 0 | 0 | 0 |
| Penmar | Penium margaritaceaum | Charophytes | Desmidiales | Peniaceae | 10.1016/j.cell.2020.04.019 | 0 | 0 | 0 |
| Petaxi | Petunia axillaris | Angiosperms | Solanales | Solanaceae | 10.1038/nplants.2016.74 | 1 | 1 | 0 |
| Phaequ | Phalaenopsis equestris | Angiosperms | Asparagales | Orchidaceae | 10.1038/nature23897 | 1 | 0 | 0 |
| Phavul | Phaseolus vulgaris | Angiosperms | Fabales | Fabaceae | 10.1038/ng.3008 | 1 | 1 | 1 |
| Phypat | Physcomitrium patens | Mosses | Funariales | Funariaceae | 10.1111/tpj.13801 | 0 | 0 | 0 |
| Picabi | Picea abies | Gymnosperms | Pinales | Pinaceae | 10.1038/nature12211 | 0 | 0 | 0 |

| | | | | | | | | |
|---|---|---|---|---|---|---|---|---|
| Picgla | Picea glauca | Gymnosperms | Pinales | Pinaceae | 10.1093/bioinformatics/btt178 | 0 | 0 | 0 |
| Picsit | Picea sitchensis | Gymnosperms | Pinales | Pinaceae | Gymno-plaza | 0 | 0 | 0 |
| PicspRCC4223 | Picochlorum sp. RCC4223 | Chlorophytes | Chlorellales | Incertae sedis | ORCAE | 0 | 0 | 0 |
| Pinpin | Pinus pinaster | Gymnosperms | Pinales | Pinaceae | Gymno-plaza | 0 | 0 | 0 |
| Pinsyl | Pinus sylvatica | Gymnosperms | Pinales | Pinaceae | 10.1038/nature12211 | 0 | 0 | 0 |
| Pintae | Pinus taeda | Gymnosperms | Pinales | Pinaceae | 10.15334/genetics.113.159715 | 0 | 0 | 0 |
| Pissat | Pisum sativum | Angiosperms | Fabales | Fabaceae | 10.1038/s41588-019-0480-1 | 1 | 1 | 1 |
| Popalb | Populus alba | Angiosperms | Malpighiales | Salicaceae | NCBI | 1 | 1 | 0 |
| Popeup | Populus euphratica | Angiosperms | Malpighiales | Salicaceae | NCBI | 1 | 1 | 0 |
| Poptri | Populus trichocarpa | Angiosperms | Malpighiales | Salicaceae | 10.1126/science.1128691 | 1 | 1 | 0 |
| Porpur | Porphyridium purpureum | SAR | Porphyridiales | Porphyridiaceae | 10.1038/ncomms2931 | 0 | 0 | 0 |
| Porumb | Porphyra umbilicalis | SAR | Bangiales | Bangiaceae | 10.1073/pnas.1703088114 | 0 | 0 | 0 |
| Potmic | Potentilla micrantha | Angiosperms | Rosales | Rosaceae | 10.1093/gigascience/giy010 | 1 | 1 | 0 |
| Pracol | Prasinoderma coloniale | Prasinodermophytes | Prasinodermales | Prasinodermophyceae | 10.1038/s41559-020-1221-7 | 0 | 0 | 0 |
| Proalb | Prosopis alba | Angiosperms | Fabales | Fabaceae | NCBI | 1 | 1 | 1 |
| Procyn | Protea cynaroides | Angiosperms | Proteales | Proteaceae | 10.1111/tpj.16044 | 0 | 0 | 0 |
| Prowic | Prototheca wickerhamii | Chlorophytes | Chlorellales | Chlorellaceae | NCBI | 0 | 0 | 0 |
| Pruavi | Prunus avium | Angiosperms | Rosales | Rosaceae | 10.1093/dnares/dsx020 | 1 | 1 | 0 |
| Prudul | Prunus dulcis | Angiosperms | Rosales | Rosaceae | 10.1111/tpj.14538 | 1 | 1 | 0 |
| Prumum | Prunus mume | Angiosperms | Rosales | Rosaceae | NCBI | 1 | 1 | 0 |
| Pruper | Prunus persica | Angiosperms | Rosales | Rosaceae | 10.1038/ng.2586 | 1 | 1 | 0 |
| Pruyed | Prunus yedoensis | Angiosperms | Rosales | Rosaceae | NCBI | 1 | 1 | 0 |

| | | | | | | | | |
|---|---|---|---|---|---|---|---|---|
| Psemen | Pseudotsuga menziesii | Gymnosperms | Pinales | Pinaceae | 10.1534/g3.117.300078 | 0 | 0 | 0 |
| Pyrbet | Pyrus betulifolia | Angiosperms | Rosales | Rosaceae | NCBI | 1 | 1 | 0 |
| Pyrbre | Pyrus x bretschneideri | Angiosperms | Rosales | Rosaceae | NCBI | 1 | 1 | 0 |
| Pyrcom | Pyrus communis | Angiosperms | Rosales | Rosaceae | 10.1371/journal.pone.0092644 | 1 | 1 | 0 |
| Pyryez | Pyropia yezoensis | SAR | Bangiales | Bangiaceae | 10.1371/journal.pone.0057122 | 0 | 0 | 0 |
| Quelob | Quercus lobata | Angiosperms | Fagales | Fagaceae | NCBI | 1 | 1 | 0 |
| Querob | Quercus robur | Angiosperms | Fagales | Fagaceae | 10.1111/1755-0998.12425 | 1 | 1 | 0 |
| Rharub | Rhamnella rubrinervis | Angiosperms | Rosales | Rhamnaceae | NCBI | 1 | 1 | 0 |
| Rhoirr | Rhododendron irroratum | Angiosperms | Ericales | Ericaceae | 10.3389/fpls.2023.1123707 | 1 | 0 | 0 |
| Rhosim | Rhododendron simsii | Angiosperms | Ericales | Ericaceae | 10.1038/s41467-020-18771-4 | 1 | 0 | 0 |
| Rhowil | Rhododendron williamsianum | Angiosperms | Ericales | Ericaceae | 10.1093/gbe/evz245 | 1 | 0 | 0 |
| Riccom | Ricinus communis | Angiosperms | Malpighiales | Euphorbiaceae | 10.1038/nbt.1674 | 1 | 1 | 0 |
| Roraqu | Rorripa aquatica | Angiosperms | Brassicales | Brassicaceae | 10.1101/2022.06.06.494894 | 0 | 0 | 0 |
| Roschi | Rosa chinensis | Angiosperms | Rosales | Rosaceae | 10.1038/s41588-018-0110-3 | 1 | 1 | 0 |
| Rubocc | Rubus occidentalis | Angiosperms | Rosales | Rosaceae | 10.1111/tpj.13215 | 1 | 1 | 0 |
| Salcuc | Salvinia cucullata | Monilophytes | Salviniales | Salviniaceae | 10.1038/s41477-018-0188-9 | 0 | 0 | 0 |
| Sclbir | Sclerocarya birrea | Angiosperms | Sapindales | Anarcadiaceae | 10.1093/gigascience/giy152 | 1 | 1 | 0 |
| Sellep | Selaginella lepidophylla | Lycophytes | Selaginellales | Selaginellaceae | 10.1038/s41467-017-02546-5 | 1 | 1 | 0 |
| Selmoe | Selaginella moellendorffii | Lycophytes | Selaginellales | Selaginellaceae | 10.1126/science.1203810 | 1 | 1 | 0 |
| Sentor | Senna tora | Angiosperms | Fabales | Fabaceae | 10.1038/s41467-021-21987-7 | 1 | 1 | 0 |
| Setita | Setaria italica | Angiosperms | Poales | Poaceae | 10.1038/nbt.2196 | 1 | 1 | 0 |

| | | | | | | | | |
|---|---|---|---|---|---|---|---|---|
| Sollyc | Solanum lycopersicum | Angiosperms | Solanales | Solanaceae | 10.1038/nature11119 | 1 | 1 | 0 |
| Solpen | Solanum pennellii | Angiosperms | Solanales | Solanaceae | 10.1038/nature11119 | 1 | 1 | 0 |
| Sorbic | Sorghum bicolor | Angiosperms | Poales | Poaceae | 10.1111/tpj.13781 | 1 | 1 | 0 |
| Spasub | Spatholobus suberectus | Angiosperms | Fabales | Fabaceae | NCBI | 1 | 1 | 1 |
| Sphfal | Sphagnum fallax | Mosses | Sphagnales | Sphagnaceae | Phytozome | 0 | 0 | 0 |
| Spimus | Spirogloea muscicola | Charophytes | Zygnemales | Zygnematophyceae | 10.1016/j.cell.2019.10.019 | 0 | 0 | 0 |
| Spiole | Spinacia oleracea | Angiosperms | Caryophyllales | Amaranthaceae | bvseq.molgen.mpg.de | 0 | 0 | 0 |
| Spipol | Spirodela polyrhiza | Angiosperms | Alismatales | Araceae | 10.1038/ncomms4311 | 0 | 0 | 0 |
| SymirrSAG2036 | Symbiochloris irregularis SAG2036 | Chlorophytes | Trebouxiales | Trebouxiaceae | 10.1101/2022.01.06.475074 | 0 | 0 | 0 |
| Tarhas | Tarenaya hassleriana | Angiosperms | Brassicales | Cleomaceae | 10.1105/tpc.113.113480 | 0 | 0 | 0 |
| Thecac | Theobrama cacao | Angiosperms | Malvales | Malvaceae | 10.1186/gb-2013-14-6-r53 | 1 | 1 | 0 |
| TregelLA000220 | Trebouxia gelatinosa | Chlorophytes | Trebouxiales | Trebouxiaceae | 10.1007/s11103-016-0468-5 | 0 | 0 | 0 |
| Treori | Trema orientalis | Angiosperms | Rosales | Cannabaceae | 10.1073/pnas.1721395115 | 1 | 1 | 0 |
| TrespOTU1 | Trebouxia sp. OTU1 generalist lineage | Chlorophytes | Trebouxiales | Trebouxiaceae | 10.1101/2022.01.06.475074 | 0 | 0 | 0 |
| TrespOTU3 | Trebouxia sp. OTU3 warm lineage | Chlorophytes | Trebouxiales | Trebouxiaceae | 10.1101/2022.01.06.475074 | 0 | 0 | 0 |
| TrespOTU5 | Trebouxia sp. OTU3 cold lineage | Chlorophytes | Trebouxiales | Trebouxiaceae | 10.1101/2022.01.06.475074 | 0 | 0 | 0 |
| TrespTZW2008 | Trebouxia sp. TZW2008 | Chlorophytes | Trebouxiales | Trebouxiaceae | 10.1186/s12864-020-07086-9 | 0 | 0 | 0 |
| Triaes | Triticum aestivum IWGSC_v1.1 | Angiosperms | Poales | Poaceae | 10.1093/gigascience/gix097 | 1 | 1 | 0 |

| | | | | | | | | |
|---|---|---|---|---|---|---|---|---|
| | _HC_20170706 | | | | | | | |
| Tripra | Trifolium pratense | Angiosperms | Fabales | Fabaceae | 10.1038/srep17394 | 1 | 1 | 1 |
| Trisub | Trifolium subterraneum | Angiosperms | Fabales | Fabaceae | NCBI | 1 | 1 | 1 |
| Ulvmut | Ulva mutabilis | Chlorophytes | Ulvales | Ulvaceae | ORCAE | 0 | 0 | 0 |
| UlvproYS2018 | Ulva prolifera YS-2018 | Chlorophytes | Ulvales | Ulvaceae | NCBI | 0 | 0 | 0 |
| Utrgib | Utricularia gibba | Angiosperms | Lamiales | Lamiaceae | 10.1073/pnas.1702072114 | 0 | 0 | 0 |
| Utrren | Utricularia reniformis | Angiosperms | Lamiales | Lamiaceae | 10.3390/ijms21010003 | 0 | 0 | 0 |
| Vicfab | Vicia faba | Angiosperms | Fabales | Fabaceae | 10.1038/s41586-023-05791-5 | 1 | 1 | 1 |
| Vigang | Vigna angularis | Angiosperms | Fabales | Fabaceae | 10.1038/srep080669 | 1 | 1 | 1 |
| Vigrad | Vigna radiata | Angiosperms | Fabales | Fabaceae | 10.1038/ncomms6443 | 1 | 1 | 1 |
| Vigsub | Vigna subterranea | Angiosperms | Fabales | Fabaceae | 10.1093/gigascience/giy152 | 1 | 1 | 1 |
| Vigung | Vigna unguiculata | Angiosperms | Fabales | Fabaceae | 10.1111/tpj.14349 | 1 | 1 | 1 |
| Volcar | Volvox carteri | Chlorophytes | Chlamydomonales | Volvocaceae | 10.1126/science.1188800 | 0 | 0 | 0 |
| Xervis | Xerophyta viscosa | Angiosperms | Pandanales | Velloziaceae | 10.1038/nplants.2017.38 | 1 | 1 | 0 |
| Zeamay | Zea mays PH207 | Angiosperms | Poales | Poaceae | 10.1105/tpc.16.00353 | 1 | 1 | 0 |
| Zizjuj | Ziziphus jujuba | Angiosperms | Rosales | Rhamnaceae | 10.1038/ncomms6315 | 1 | 1 | 0 |
| Zosmar | Zostera marina | Angiosperms | Alismatales | Zosteraceae | 10.1038/nature16548 | 0 | 0 | 0 |
